# Spatial and long-term temporal evolution of a marine mussel hybrid zone (*Mytilus* spp.) in SW England

**DOI:** 10.1101/2023.07.19.549755

**Authors:** Angel P. Diz, David O. F. Skibinski

## Abstract

The study of spatial and temporal changes in hybrid zones offers important insights into speciation. Earlier studies on hybrid populations of the marine mussel species *Mytilus edulis* and *M. galloprovincialis* in SW England provided evidence of admixture but were constrained by the limited number of molecular markers available. We use 57 highly ancestry-informative SNPs, most of which have been mapped genetically, to provide evidence of distinctive differences between highly admixed populations in SW England and asymmetrical introgression from *M. edulis* to *M. galloprovincialis*. We combine the genetic study with analysis of phenotypic traits of potential ecological and adaptive significance. We demonstrate that hybrid individuals have brown mantle edges unlike the white or purple in the parental species, suggesting allelic or non-allelic genomic interactions. We report differences in gonad development stage between the species consistent with a prezygotic barrier between the species. By incorporating results from publications dating back to 1980 we confirm the long-term stability of the hybrid zone consistent with higher viability of *M. galloprovincialis*. This stability coincides with a dramatic change in temperature of UK coastal waters and suggests that these hybrid populations might be resisting the effects of global warming. However, a single SNP locus associated with the notch 2 signalling transmembrane protein shows a markedly different pattern of variation to the others and might be associated with adaption of *M. galloprovincialis* to colder northern temperatures.

## Introduction

The role of hybridisation in evolution has always been a topic of great interest for evolutionary biologists. Hybridisation is closely linked to reproductive isolation and speciation. It promotes gene flow between divergent taxa which contributes to biodiversity through the formation of novel phenotypes, generating adaptive variation, and may also lead to speciation (Abbott et al. 2013; Seehausen et al. 2014). Hybrid zones are considered excellent natural scenarios for investigating the evolutionary process and understanding how genetic and environmental factors contribute to reproductive isolation (Gompert et al. 2017; Taylor & Larson 2019). Natural hybrid zones from marine organisms have been studied less than those involving terrestrial species despite hybridization in the sea being more common than previously thought (Gardner 1997). This is changing with recent advances in genomic analysis and the integration of genomic results with other research fields (Faria et al. 2021). The *Mytilus edulis* complex comprises three different closely related species in the Northern hemisphere: *Mytilus edulis* and *M. trossulus,* which inhabit temperate to cold regions, and *M. galloprovincialis,* known as the Mediterranean mussel, found in warmer regions. Hybridisation occurs on rocky shores where their distributions overlap, forming hybrid zones that feature different ecological conditions and degrees of interspecific gene flow and admixture (Skibinski et al. 1983; Koehn 1991; Bierne et al. 2003a; Riginos & Cunningham 2005). *Mytilus* hybrid zones involving different pairs of species have been observed in the North-West (*M. edulis x M. trossulus*) and North-East (*M. edulis* x *M. galloprovincialis* and *M. edulis x M. trossulus*) Atlantic coasts. In Europe, the most well studied hybrid zones are those present along the Atlantic French and South-West (SW) England coast involving *M. edulis* and *M. galloprovincialis* (Skibinski et al. 1978a, 1983; Coustau et al. 1991; Gardner & Skibinski 1988, 1990, 1991b; Daguin et al. 2001; Hilbish et al. 2002; Bierne et al. 2003a; Simon et al. 2021).

There is much evidence of morphological, anatomical, physiological and genetic differences between *Mytilus edulis* and *M. galloprovincialis* (Lewis & Seed 1969; Seed 1971; Skibinski et al. 1983; Koehn 1991; McDonald et al. 1991; Gardner 1992; Gosling 1992; Hilbish et al. 1994). Earlier studies have drawn conclusions about species-specific phenotypic differences by analysing wild parental and putative hybrids. A limitation of these studies was the lack of precision in determining the genetic status of parental and hybrid mussels, due to the limited number of molecular markers available. This highlights the need for further analyses with a higher number of ancestry-informative molecular markers.

Reproductive barriers help preserve the genetic and physiological differences between species. Additionally, in *Mytilus* species there are differences in habitat preference (Gosling & McGrath 1990; Bierne et al. 2002b, 2003b) and assortative fertilisation (Bierne et al. 2002a). The study of reproductive timing (spawning peaks and asynchrony) has received attention as a prezygotic mechanism of reproductive isolation. Analysis of the gonad index or histological tests provided evidence of reproductive cycle differences in *M. edulis* and *M. galloprovincialis* living in sympatry. However, these results were variable between different sites and seasons with substantial spawning overlap (Seed 1971; Gardner 1992; Secor et al. 2001; Gilg & Hilbish 2003c; Bignell et al. 2008).

The study of hybrid zones may be of particular value in gauging the effects of climate change and global warming (Chunco 2014; Taylor et al. 2015). However, studies of the temporal genetic variability of marine hybrid zones are rare (Gardner 1997). The northern expansion of *M. galloprovincialis* might be limited by cold winter temperatures affecting reproductive development (Fly et al. 2015). Thus the study of the evolution of mussel hybrid zones at geographical and temporal scales using genetic and reproductive analysis might provide valuable information on the ecological response to global warming.

Population genetic studies of natural populations have progressed from the analysis of allozymes (1970s) to mtDNA (1980s), microsatellites (1990s), SNPs (2000s) to whole genomes (Allendorf 2017). In the study of *Mytilus* hybrid zones in SW England early work using 5-10 ancestry-informative allozyme markers was followed by mtDNA analysis. The use of a highly-discriminant nuclear marker (Glu-5) was introduced later. The use of a larger number of genetic markers would increase the power to better characterise individuals genetically. In *Mytilus* species ancestry-informative SNP markers have been developed recently and used to investigate the population genetics of several *Mytilus* hybrid zones and other populations showing variable levels of admixture and introgression (Zbawicka et al. 2012, 2014; Fraïsse et al 2016; Mathiesen et al. 2017; Simon et al. 2018, 2020, 2021; Vendrami et al 2020; Wenne et al. 2016, 2020). However, these techniques have not yet been applied to populations in the SW England hybrid zone.

In the present study, we use 57 ancestry-informative SNP markers obtained from previous whole genomic and transcriptomic studies in *Mytilus* spp., most of which have been mapped genetically, to analyse interbreeding and admixture between *M. edulis* and *M. galloprovincialis*. We demonstrate differences between hybrid populations, and provide support for two distinct hybrid zone regions in SW England. The results suggest asymmetrical gene flow from *M. edulis* into *M. galloprovincialis*. Evidence for stable size and age dependent genetic variation over a period of around 35 years supports the long-term stability of the zone despite warming sea temperatures. One SNP locus showed a pattern of geographic variation markedly different from the others and might reflect local adaptation.

## Material and Methods

### Sampling of mussels, tissue collection and phenotypic traits

Marine mussels (*Mytilus* spp.) were collected in March 2016 from the intertidal rocky shore in Southwest (UK) England for six populations of *Mytilus edulis* and *Mytilus galloprovincialis* including localities where these species live in sympatry and hybridise (Skibinski et al. 1978a, 1983) (**Figure 1**). The populations (with two letter abbreviations) are Croyde (CR), Bude (BD), Padstow (PD), Carlyon Bay (CB), Whitsand Bay (WB) and Porthcurno (PC). Two of these, Bude and Padstow, are considered as regional reference populations for *M. galloprovincialis*. The other four are mixed populations of both species. Samples were also taken from two allopatric reference populations, Swansea (SW) in South Wales (*M. edulis*) and Vigo (VG) in Spain (*M. galloprovincialis*). Mussels with sizes ranging from 20 to 40 mm were transported to the marine station at the University of Vigo, Spain (ECIMAT, CIM-UVigo) for processing. Mussels were sorted according to shell length, into small (20–29 mm) and large (30–40 mm) classes. For each population, 60 individual mussels (50% for each class) were taken randomly. Mantle edge tissue was dissected from each mussel and preserved in 95% ethanol for DNA extraction and genetic analyses. Gonad tissue was collected for histological analysis.

**Figure 1:**
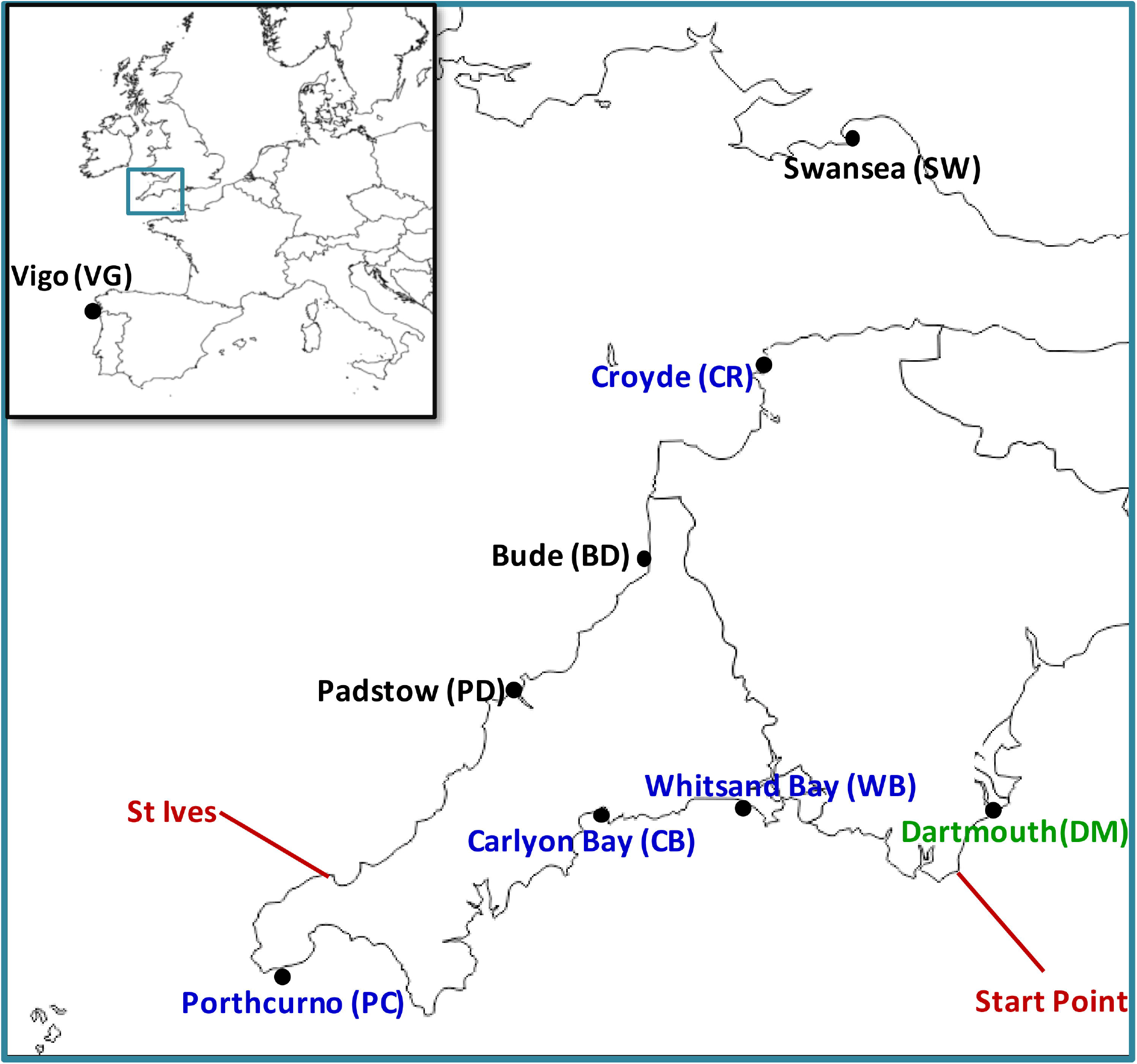
Locations of the populations sampled in 2016. GPS coordinates are as follows: Swansea, South Wales (SW): 51°34′04′′N/03°58′35′′W, Croyde (CR): 51°08′01′′N/04°14′36′′W, Bude (BD): 50°47′44′′N/04°33′28′′W, Padstow (PD): 50°33′43′′N/04°55′46′′W, Porthcurno (PC): 50°02′31′′N/05°39′02′′W, Carlyon Bay (CB): 50°20′09′′N/04°44′00′′W, Whitsand Bay (WB): 50°20′49′′N/04°15′29′′W, Vigo, North West Spain (VG): 42°11′56′′N, 08°47′22′′W. Reference population names are in black, hybrid population names in blue. An additional sample (in green) of 20 mussels (shell length ranging from 25 to 64 mm) was collected in 2018 from Dartmouth (DM): 50°20′35′′N/03°34′20′′W and analysed later for the same 57 SNP markers. Two localities which bound a hybrid zone in the south of SW England are shown in red.

Gonad development stage was assigned to seven categories (0-6) using a histological test based on hematoxylin-eosin stain (Seed 1969), with the classification scheme of Kenchington et al. (2020). Stages 0-3 do not exhibit mature gametes and were pooled together for analysis (abbreviated “Not Ripe”) as were stages 4-6 which exhibit mature gametes (abbreviated “Ripe”). 84.1% of mussels analysed were ripe or had spawned a few days before analysis (stages 4 to 6, including post-spawning). Sex was determined with this test and recorded as Female or Male. The results for gonad development stage and sex are given in **File S1-worksheet (ws) E**.

Mantle edge colour, gonad colour and the shell length and height of each mussel were also recorded and are given in **File S1-wsE**. Mantle colour was assigned as Purple, Brown, or White/Straw (abbreviated “White”), and gonad colour as deep orange (abbreviated “Orange”), intermediate, or Pale Orange/White (abbreviated “White”) (**File S1-wsD**).

### DNA extraction and genotyping

DNA was extracted from mantle edge tissue (∼ 30 mg) using the EZNA ® Mollusc DNA kit (Omega Bio-tek, USA) following manufacturer’s instructions. DNA samples were quantified and quality-checked (A260/A280 absorbance ratio) using a BioDrop µLITE (Biodrop, Cambridge, UK), standardized to 40 ng/µL and stored at −20°C. The Glu-5 marker was amplified by PCR using primers (Me15: 5’-CCA GTA TAC AAA CCT GTG AAG A, Me16: 5’-TGT TGT CTT AAT AGG TTT GTA AGA) and conditions described by Inoue et al. (1995). For genotyping, PCR products together with the NZYDNA ladder V (NZYtech, Lisbon, Portugal) were separated in 2% agarose gels supplemented with 2.5 µL of SYBR Safe dye at 90 volts for 40 minutes. Individuals were genotyped for two alleles: 126 (G) and 180 (E) base pairs long. The G or E alleles are at high frequency (>95%) in the reference populations *Mytilus galloprovincialis* and *M. edulis*respectively, used in this study.

SNP genotyping was carried out by LGC genomics (Hoddesdon, UK) following KASP™ (Kompetitive Allele Specific PCR; Semagn et al. 2014) for 57 SNP markers which show high F_ST_ values between allopatric reference populations from *Mytilus edulis* and *M. galloprovincialis* (Simon et al. 2018, 2020, 2021). They were developed from targeted enrichment sequencing data (Fraïsse et al. 2016). The alleles at each locus were recoded as G or E from the original DNA base assignation according to whether they were at higher frequency in *M. galloprovincialis* or *M. edulis* respectively. The sequences amplified by KASP™ which contained the 57 SNPs were used for blastX against the nr NCBI protein database. This was done to assess whether the KASP™ sequences were located in coding or non-coding regions, and whether the SNP substitutions were synonymous or non-synonymous. The KASP™ sequences and the blastX results are given in **File S1-wsA** and **wsB**.

### Data analysis

Raw genotypic data for each SNP marker and for Glu-5, together with information for the phenotypic traits studied are provided in **File S1-wsC** and **wsE**. The totals for the EE, EG, and GG genotypes counted across the 57 SNP loci and the proportion of SNP loci for which an individual is heterozygous (Ho) are given. Genotype and allele frequencies are given for each SNP locus for each population in **File S1-**from **wsF** to **wsM**. These worksheets also give the observed and expected heterozygosity, the F_IS_ statistic, and p-value for test for Hardy-Weinberg equilibrium. Values are also given separately for mussels of the two different shell length classes. F_IS_ and p-values were calculated in Genepop vs.4.7.5 (Rousset 2008) using the method of Weir and Cockerham (1984).

Contingency analyses were carried out in SPSS vs.28 using Fisher’s Exact test with the Monte Carlo method based on 100,000 sampled tables. The significance of individual cells was determined by converting the adjusted standardised residual to a two-tail p-value using the cumulative distribution function (CDF) (function norm.s.dist in Excel). As a measure of association, Cramer’s V test was calculated for each contingency table.

STRUCTURE vs.2.3.4 program (Pritchard et al. 2000) was used to estimate the ancestral probability for each individual mussel (Q-value). This can be interpreted as a measure of *M. edulis* and *M. galloprovincialis* ancestry expressed as a proportion between 0 and 1. Run parameters were set to k=2, assuming an admixture and correlated allele frequencies model, with 100,000 burn in and 1,000,000 iteration steps. A genetic map of 36 SNP markers was constructed for *Mytilus* by Simon et al. (2020). 32 of those SNP markers were used in the present study. They comprise six groups of linked markers and a further 9 unlinked markers (**File S2-wsA**). With these markers, STRUCTURE was used to estimate the number of generations since a recent admixture event in the hybrid populations using Swansea and Bude as reference populations (POPFLAG option ON), following a linkage model (Falush et al. 2003).

NewHybrids vs.1.1 (Anderson & Thompson 2002) was used to calculate the posterior probabilities that individuals in the population samples belong to different genotypic frequency categories for two generations of interbreeding (G2). They are P0 (*M. edulis*), P1 (*M. galloprovincialis*), F1, F2, and backcrosses BC0 and BC1 and are distinguished by having different expected genotypic proportions for EE, EG, GE and GG (**File S2-wsB**). They are distinct from genealogical classes which can be named p0, p1, f1, f2, bc1 and bc2. Thus, a cross between an *M. galloprovincialis* parent (p1) and a first-generation hybrid (f1) would produce a backcross individual (bc1). It is possible that an individual belonging to a particular genealogical class (e.g., bc1) might not generate a high posterior probability to the corresponding genotypic frequency category (i.e., BC1), but to another. This may happen when the individual belongs to a genealogical class whose corresponding genotypic frequency category is not amongst those presented in the analysis. This could happen when a G2 analysis is used for individuals produced in a third generation of interbreeding. For downstream analysis, individuals were assigned to the genotypic category for which they had the highest posterior probability.

More details of the NewHybrids analysis as applied in the present study are given in **File S3**. This also presents an exploratory analysis using the genotypic frequency categories for the third generation of interbreeding (G3).

Bayescan v.2.1 (Foll & Gaggiotti 2008) was used to check for loci with outlying F_ST_ values. The program computes a locus specific component (alpha) which indicates departure from neutrality. High and low Fst (or alpha) values suggest diversifying selection or purifying selection respectively. We used prior odds for the neutral model of 10 which is the default. As the SNP loci were chosen a priori because they discriminated between *M. edulis* and *M. galloprovincialis* the analysis was applied separately only to P0 and P1 individuals from the six populations in the hybrid zone (Swansea and Vigo excluded). A test for outlier loci was also carried out using the Mahalanobis distance. The calculation was carried out in SPSS vs.28 and the distance was converted to a p-value which was converted to a q-value (Storey 2002) employing the Bioconductor vs.2.28.0 qvalue package.

## Results

### Genetic Structure of Populations

The EE, EG, and GG genotype proportions, the G allele proportion, and F_IS_ were computed for each SNP over all mussels within each population. These summary values were then averaged over all SNP loci (**Figure 2).** Values are also given separately for small and large size classes. The *M. edulis* reference population (Swansea, SW) has the lowest average G allele frequency, whereas the *M. galloprovincialis* reference populations Bude (BD), Padstow (PD) and Vigo (VG) have the highest frequencies. The other four populations Croyde (CR), Porthcurno (PC), Carlyon (CB), and Whitsand (WB) referred to as hybrid populations have more intermediate allele frequencies. All populations have an overall deficit of heterozygotes (positive F_IS_). The values for the hybrid populations are higher and highly significant. This is as expected for mixed populations of *M. edulis* and *M. galloprovincialis* which have some restriction of interbreeding. These results are in accord with those of past studies of the genetic structure in SW England (Skibinski et al. 1978a; Skibinski 1983; Gardner & Skibinski 1988; Wilhelm & Hilbish 1998; Hilbish et al. 2002).

**Figure 2:**
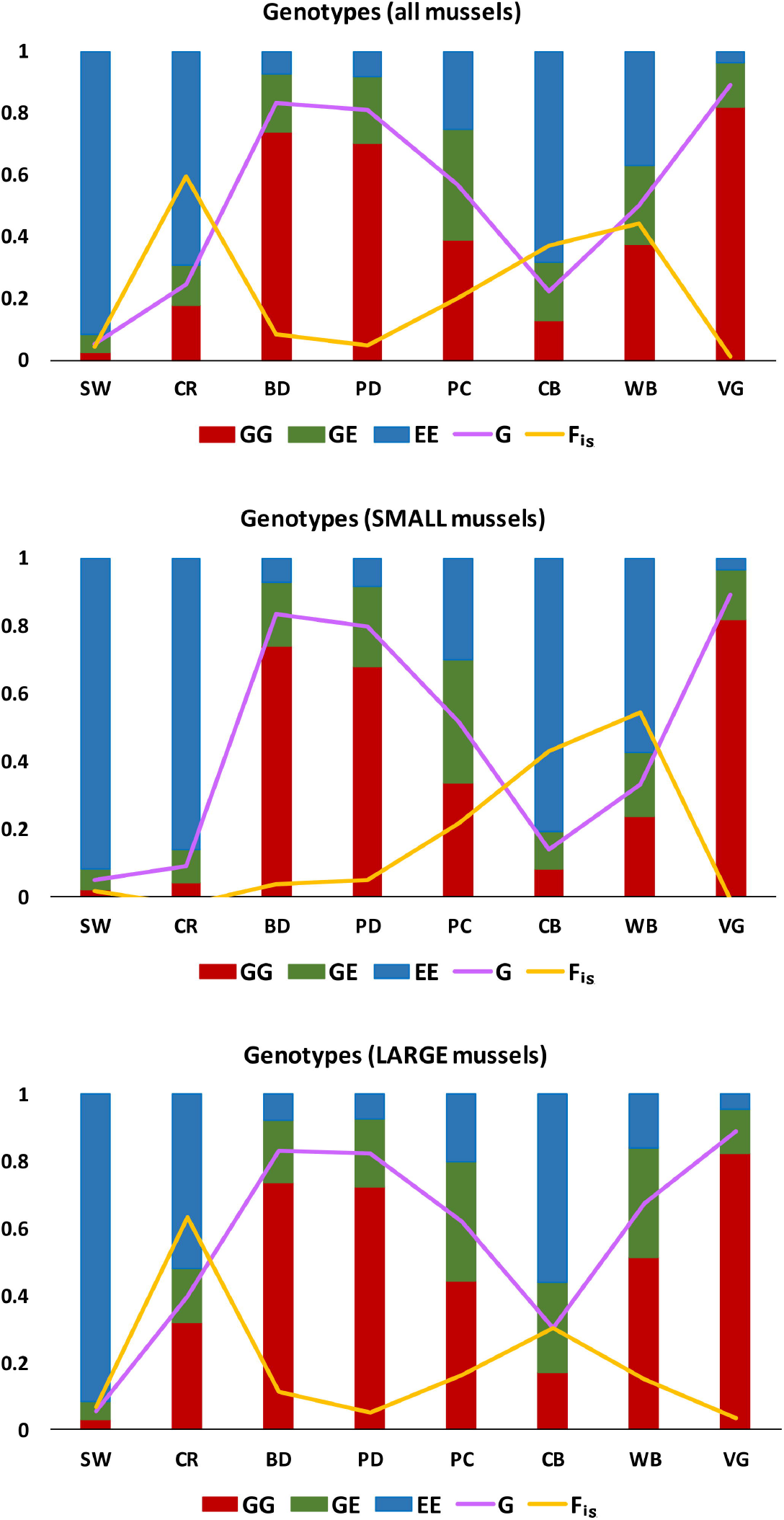
Genotype proportions (EE, EG, and GG), G allele proportion, and F_IS_ averaged over 60 individuals and 57 SNPs. Values are given for all mussels and separately for small (20-29 mm) and large (30-40 mm) mussels. The corresponding numerical values are given in **File S1-wsP**. The underlying raw data is given in **File S1-wsF** to **wsM** which also give calculated p-values for F_IS_. Population abbreviations are defined in the text.

Previous allozyme studies demonstrated a trend whereby the E allele dominates in small (and younger) mussels and the G allele in large (and older) mussels within hybrid populations (Skibinski 1983; Gardner & Skibinski 1988; Skibinski & Roderick 1991). A similar observation was made for the Glu-5 marker (Hilbish et al. 2002). This has been investigated here by comparing mussels in the two size classes. In line with the earlier studies, the G allele is greater in frequency in large mussels in the four hybrid populations (**Figure 2**). At Croyde, F_IS_ is low in small mussels and characteristic of the Swansea *M. edulis* reference population but higher in the larger mussels, characteristic of a mixed population. At Porthcurno, F_IS_ is lower in large mussels compared with Croyde indicating a different genetic structure. It is similar in value in the two size classes, consistent with a similar genetic structure characterised by some interbreeding. At Carlyon and Whitsand, F_IS_ is high in the smaller mussels, indicative of mixed populations. In the larger mussels, F_IS_ is lower indicative of greater interbreeding. These contrasts between large and small are apparent in the pattern of F_IS_ values for the individual SNP loci (**File S1-wsN**).

Scatterplots of Ho against *M. edulis* ancestry (Q-value) are shown for each population in **Figure 3**. Such “triangle” plots are informative in assessing genetic structure (Barton 2000; Fitzpatrick 2012). In reference or mixed populations of *M. edulis* and *M. galloprovincialis* without interbreeding, individuals will be clustered at low Ho to the left (e.g., Vigo, Bude and Padstow) and right (e.g., Swansea) corners of the triangle. As interbreeding proceeds, F1 individuals will appear at the apex (e.g., Croyde). The plots capture some information about deviation from Hardy-Weinberg equilibrium and about linkage disequilibrium. Thus, Croyde having many individuals to the left and right has a deficit of heterozygotes (high F_IS_, **Figure 2**) and strong linkage disequilibrium arising from the mixture of two genetically different types.

**Figure 3:**
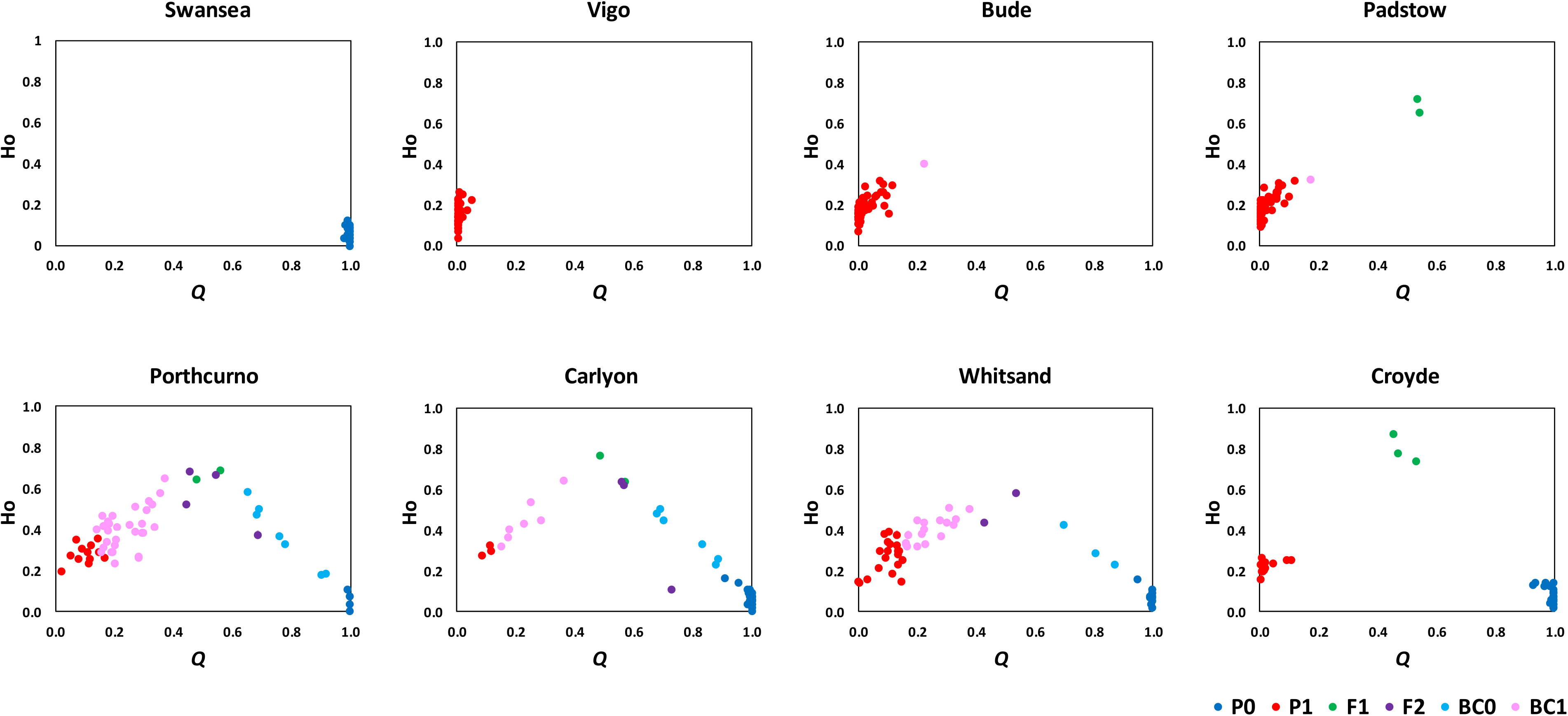
Plots of the proportion of SNP loci for which an individual is heterozygous (Ho) against Q-value (*M. edulis* ancestry) for samples from eight populations. The plotted points for individual mussels are coloured according to the NewHybrid G2 analysis genotypic frequency categories. The source data is also given in **File S1-wsE**.

The plotted points coloured according to the G2 NewHybrids genotypic frequency categories are in different regions of the triangle. This is particularly evident when the data from the eight population samples are combined (**File S2-wsJ**). The groups of points are positioned in accord with expectation given that there is some correspondence between genotypic frequency category and genealogical class. BC0 and BC1 are to the right and left of the triangle. F2 tend to have lower Ho than F1 and are more scattered as expected from segregation in f1 individuals. The extended distribution within the BC0 and BC1 classes can also be attributed to segregation in parents producing a range of values differing in both Q and Ho. Individuals assigned to the categories BC0, BC1, and F2 might be heterogeneous in genealogical class and the result of more than two generations of interbreeding.

Individuals in the Swansea and Vigo reference populations have Q values within a narrow range consistent with no history of interbreeding and admixture. The *M. galloprovincialis* populations Bude and Padstow have a few individuals assigned to the F1 and BC1 categories suggesting some interbreeding and admixture. In addition, the individuals assigned to P1 have a wider range of Q values than at Vigo. One possible explanation is that *M. galloprovincialis* in SW England have a genome with a higher proportion of genes that derive from *M. edulis*. A second possibility is that some of these individuals are genealogically the result of third or later generations of interbreeding. It is difficult to decide between these possibilities with the present data, but some relevant analysis is given in **File S3**. Croyde on the north coast of SW England (**Figure 1**) shows a more limited level of interbreeding and admixture than Porthcurno, Carlyon and Whitsand on the south coast which show a wide spread of Q and Ho values.

### Correlations Between SNP Loci

Early work with the two allozyme loci Est-D and Lap-1 used two four-point correlation coefficients (R1 and R2) computed from the observed numbers of the nine possible two locus genotypes formed from diagnostic E and G alleles (Skibinski et al. 1978a,b; Skibinski et al. 1983). R1 measures the association of the four two locus homozygote genotypes. With no admixing, R1 is high but declines as assortment in non-parental individuals generates EE GG and GG EE individuals. R2 measures the relative frequency of backcross (EE EG, EG EE, GG EG, and EG GG) compared with F1 (EG EG) and other genotypes. If interbreeding occurs to produce F1 individuals, R2 is initially high but as backcrossing occurs R2 declines. The computation of R1 and R2 are illustrated in **File S2-wsK** with examples using selected pairs of SNPs. Taking advantage of the linkage map (**File S2-wsA**), R1 and R2 were computed for the same 27 pairs of SNPs for each hybrid population, focusing on those which had low pairwise genetic identify (Nei 1972) and covering a range of genetic linkage (cM) from close to zero to unlinked. R1, R2 and map distance (cM) are plotted for the 27 pairs in **Figure 4**.

**Figure 4:**
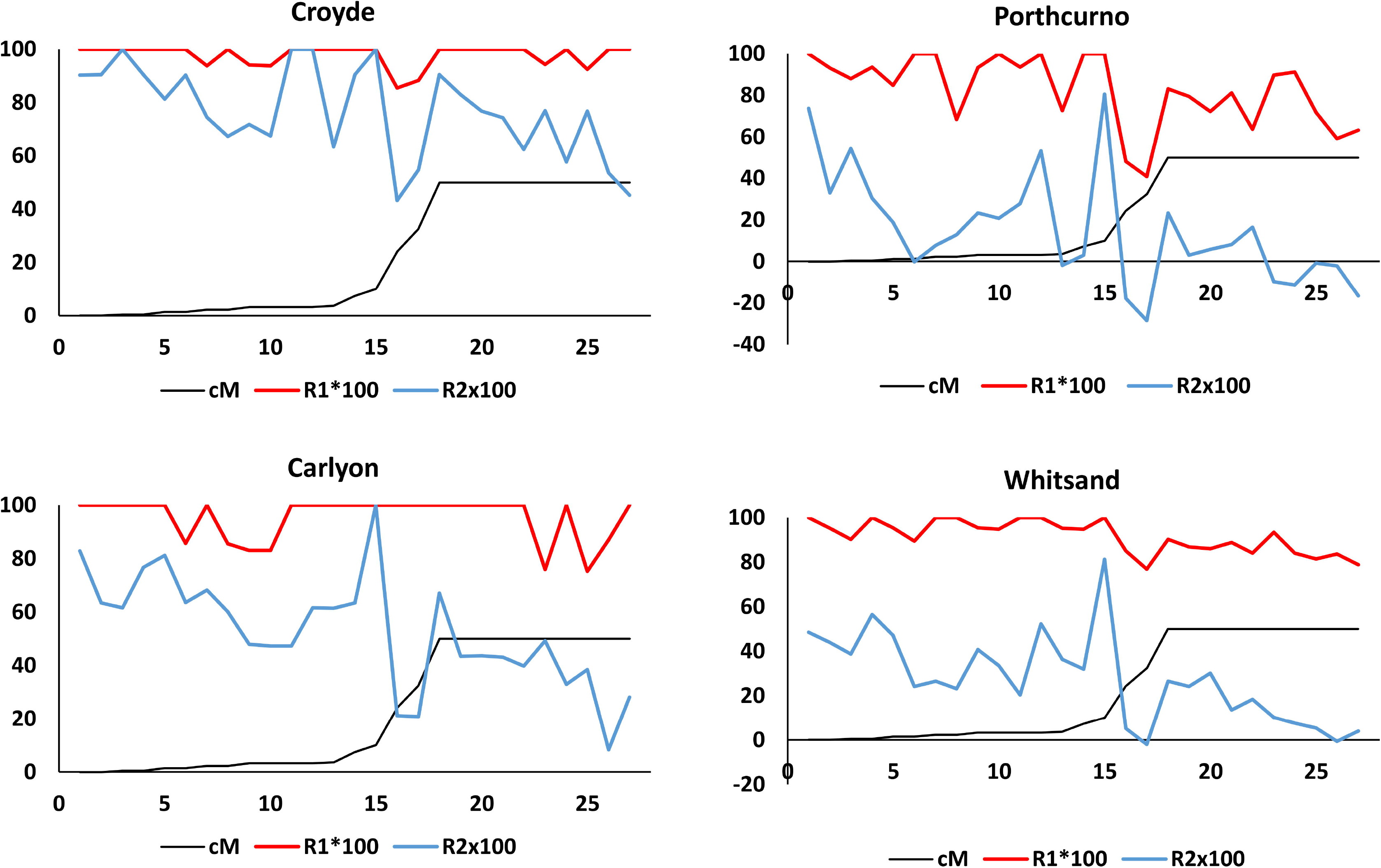
Map distance (cM) between SNP markers on a genetic map (**File S2-wsA**) (Y axis) and four-point correlation coefficients (R1 and R2) (Y-axis) plotted for 27 pairs of SNP loci ranked left to right from low to high map distance (X-axis) for four hybrid populations.

There are fluctuations in the plotted values, perhaps due to sampling error and in addition the SNP loci are not perfectly diagnostic. However, there is a general decline in both R1 and R2 as the map distance increases. This is expected as higher recombination will increase the chance of generating genotypes which are not parental or F1. The decline starts at around 30cM. The lower value of R2 for Porthcurno, Carlyon and Whitsand compared with Croyde indicates a higher level of backcrossing and is in line with the triangle plots (**Figure 3**). At higher map distance, Croyde and Carlyon maintain higher values of R1 than Porthcurno and Whitsand. Thus, although backcross individuals are frequently generated at Carlyon (low R2 and unlike Croyde where R2 is higher) admixture is not sufficient to generate EE GG and GG EE individuals to give lower R1 as in Porthcurno and Whitsand.

STRUCTURE was also used to estimate the number of generations since admixture based on estimates of recombination frequency from the genetic map of some of the SNPs. In this analysis the UK populations Bude and Swansea were used as reference populations for *M. galloprovincialis* and *M. edulis* respectively. The estimates for number of generations are 0.2 (Croyde), 2.0 (Porthcurno), 2.3 (Carlyon) and 4.3 (Whitsand).

### Population and Size Variation in NewHybrid G2 Categories

Summary histograms for the six genotypic categories are shown in **Figure 5**. For comparative analysis, three population x category contingency tests were carried out for all mussels, and small and large size classes for the four hybrid populations. First, the relative frequencies of P0 and P1 were compared. Second, the pools (P0+P1) and (F1+F2+BC0+BC1) were compared. Third, the four hybrid categories hybrid F1, F2, BC0, BC1 were compared. Detailed results are given in **File S2-wsH**. In eight of the nine tests the overall contingency tables are significant indicating differences between hybrid populations. In accord with **Figure 3** and **Figure 5**, P0 and P1 differ in frequency between the hybrid populations, and there is a significant excess of F1 at Croyde and deficit of BC1 at Croyde and Carlyon. The tendency for BC1 to increase in frequency from Carlyon to Whitsand to Porthcurno (**Figure 5**) appears to match a decline in the value of R2 (**Figure 4**) particularly at high map distance.

**Figure 5:**
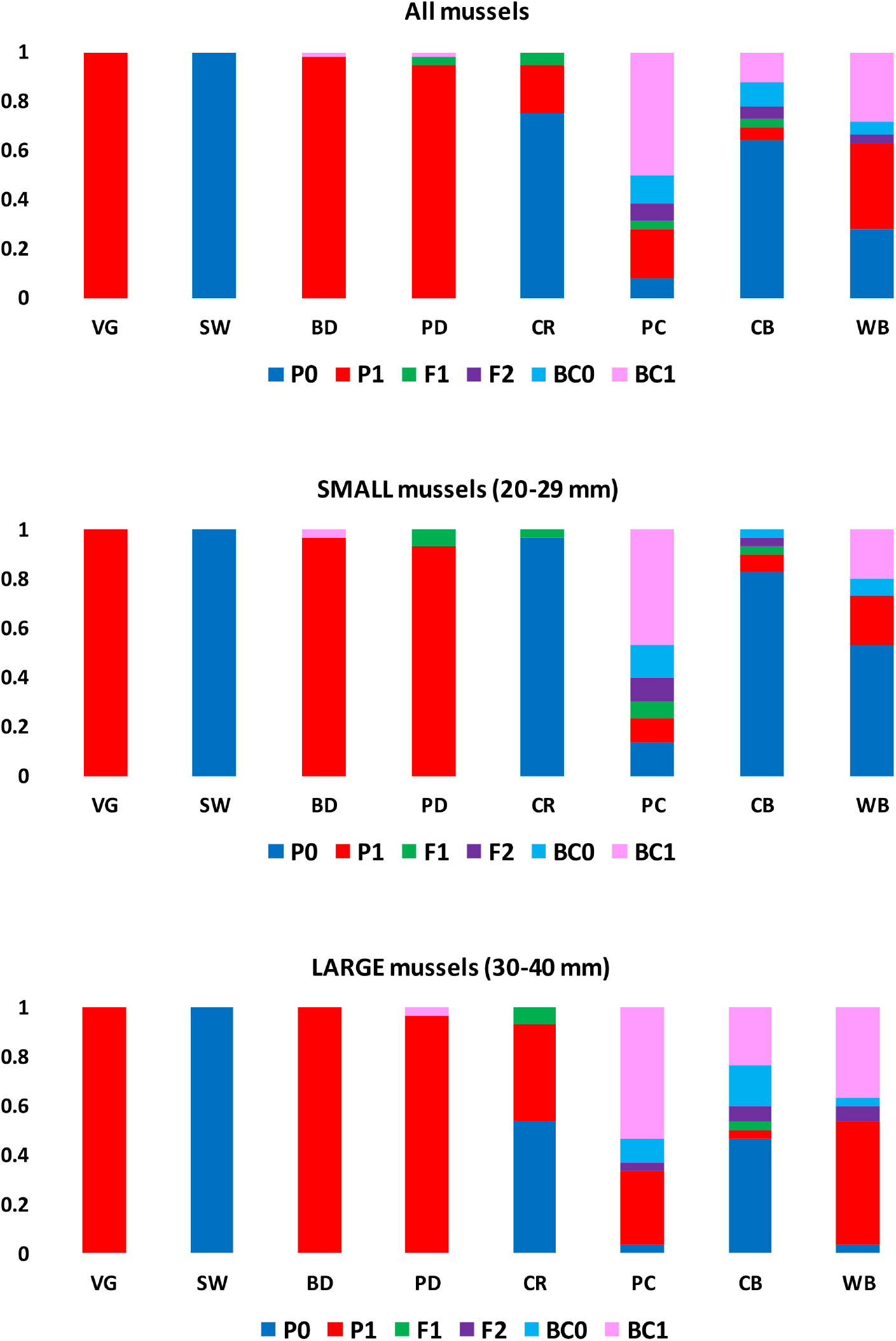
NewHybrids genotypic category assignations (G2) summed over mussels within populations for all mussels and separately for small and large mussels.

There are significant differences in the genotypic category frequency distributions between large and small size classes for the individual hybrid populations (**File S2-wsI**). In accord with **Figure 5**, P0 and P1 show a significant excess in small and large mussels respectively at Croyde, Carlyon and Whitsand but not Porthcurno, and BC1 has a significantly higher frequency in large mussels at Carlyon and in all mussels pooled. **Figure 3** and **Figure 5** both suggest a higher frequency of BC1 than BC0 individuals. Over all mussels, the ratio of BC1 to BC0 is 54 to 16. This is significant in a binomial test against a 1:1 ratio (p=0.000).

### Outlier SNP Loci

The BayeScan analysis of the 57 SNP loci plus Glu-5 for the six populations within the hybrid zone reveal no outlying loci for P0 individuals. For P1 individuals there are two outlier loci, SNP18 and SNP32 with high F_ST_ (**File S2-wsL**). These two loci have positive alpha values consistent with diversifying selection. Mahalanobis distance revealed SNP18 alone as being a significant outlier (**File S2-wsL**). Plots of G allele proportions for the P1 individuals for the six populations within the hybrid zone region demonstrate the contrast between SNP18 and the other SNP loci (**File S2-wsL**). The DNA marker Glu-5 shows a pattern across populations which correlates with the mean G allele proportion across the SNP loci, with coefficient of determination (R^2^) of 0.994 (**File S1-wsQ**). The geographic context of variation at SNP18 in SW England is given in **Figure 6**, where the E and G allele proportions are shown for the categories P0 and P1. The E allele proportion is high in P0 individuals in both the north and south coasts of SW England. However, in P1 individuals the G allele though having the expected high frequency in the southern populations Porthcurno, Carlyon and Whitsand, occurs at a low frequency in the northern populations Padstow, Bude and Croyde. Two possibilities are that directional selection has acted against the G allele and in favour of the E allele in these northern populations, or that there has been local introgression of the E allele into these populations.

**Figure 6:**
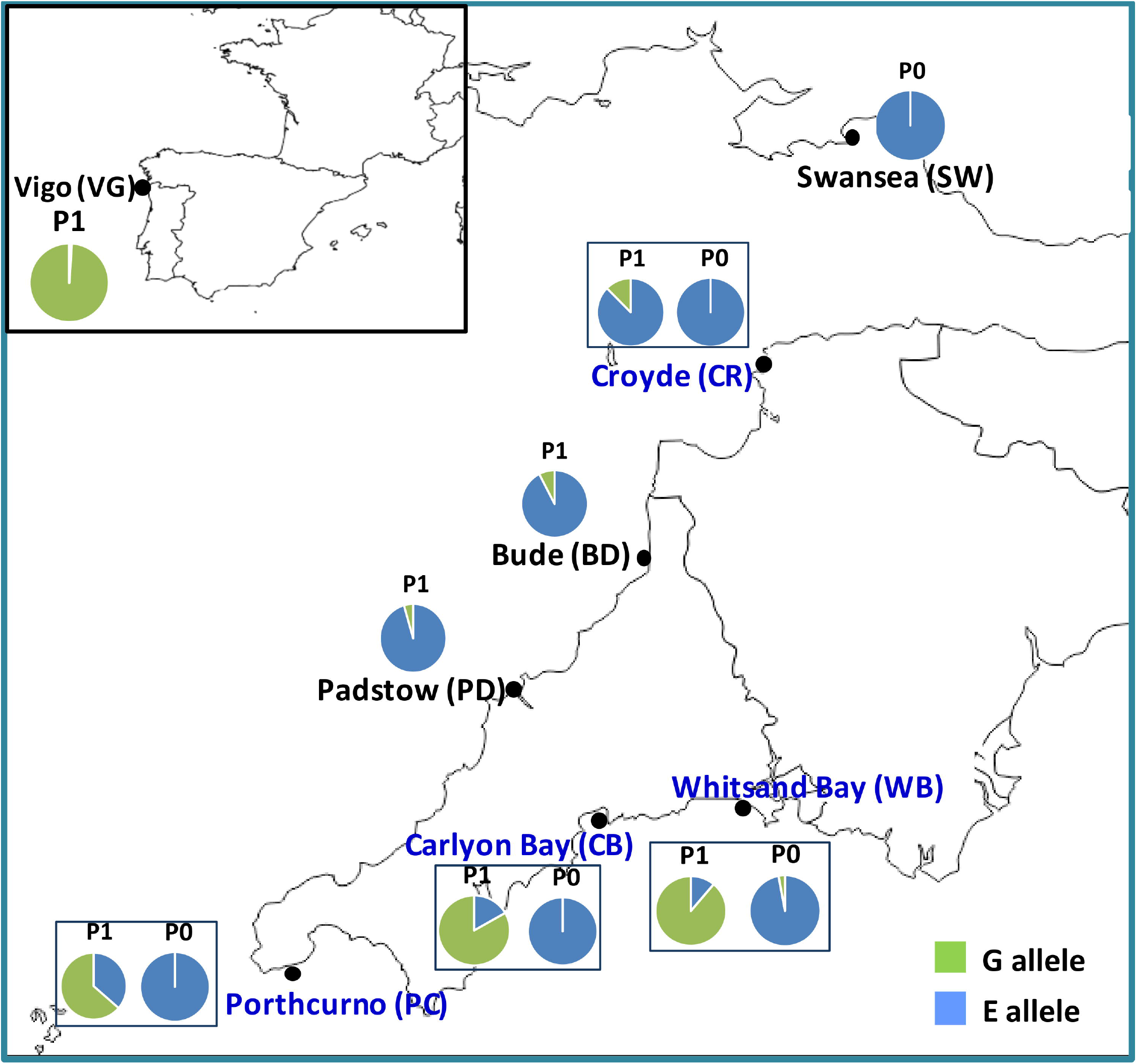
Map showing values for the proportion of E allele and G allele in P0 and P1 individuals in the eight populations for SNP18.

The SNP18 locus thus represents a good candidate to investigate the nature of any selection that might be acting on it. BlastX with the sequence within which the SNP18 polymorphism occurs gave best hits for the Notch 2 signalling transmembrane protein from *Mytilus*. The polymorphism occurs in a codon where a cytosine in *M. edulis* is substituted by an adenine in *M. galloprovincialis* associated with a nonsynonymous change from proline to glutamine. The Gene Ontology term associated with notch (GO: 0007219) links the “Notch signaling pathway” to “a series of molecular signals initiated by the binding of an extracellular ligand to the receptor Notch on the surface of a target cell, and ending with regulation of a downstream cellular process, e.g., transcription”.

### Associations Involving Phenotypic Characters

Contingency analyses were used to examine the association of the phenotypic characters (sex, mantle edge and gonad colour and gonad developmental stage) with populations and G2 genotypic categories. The main findings are reported here (**Figure 7**) but a more detailed analysis is described in **File S3** with further detailed results in **File S4**. Because sex is important in relation to doubly uniparental inheritance of mitochondrial DNA (DUI) in *Mytilus* (Fisher & Skibinski 1990; Skibinski et al. 1994a,b; Zouros et al. 1994a,b), the relationship between sex and the other phenotypic characters was also analysed.

**Figure 7:**
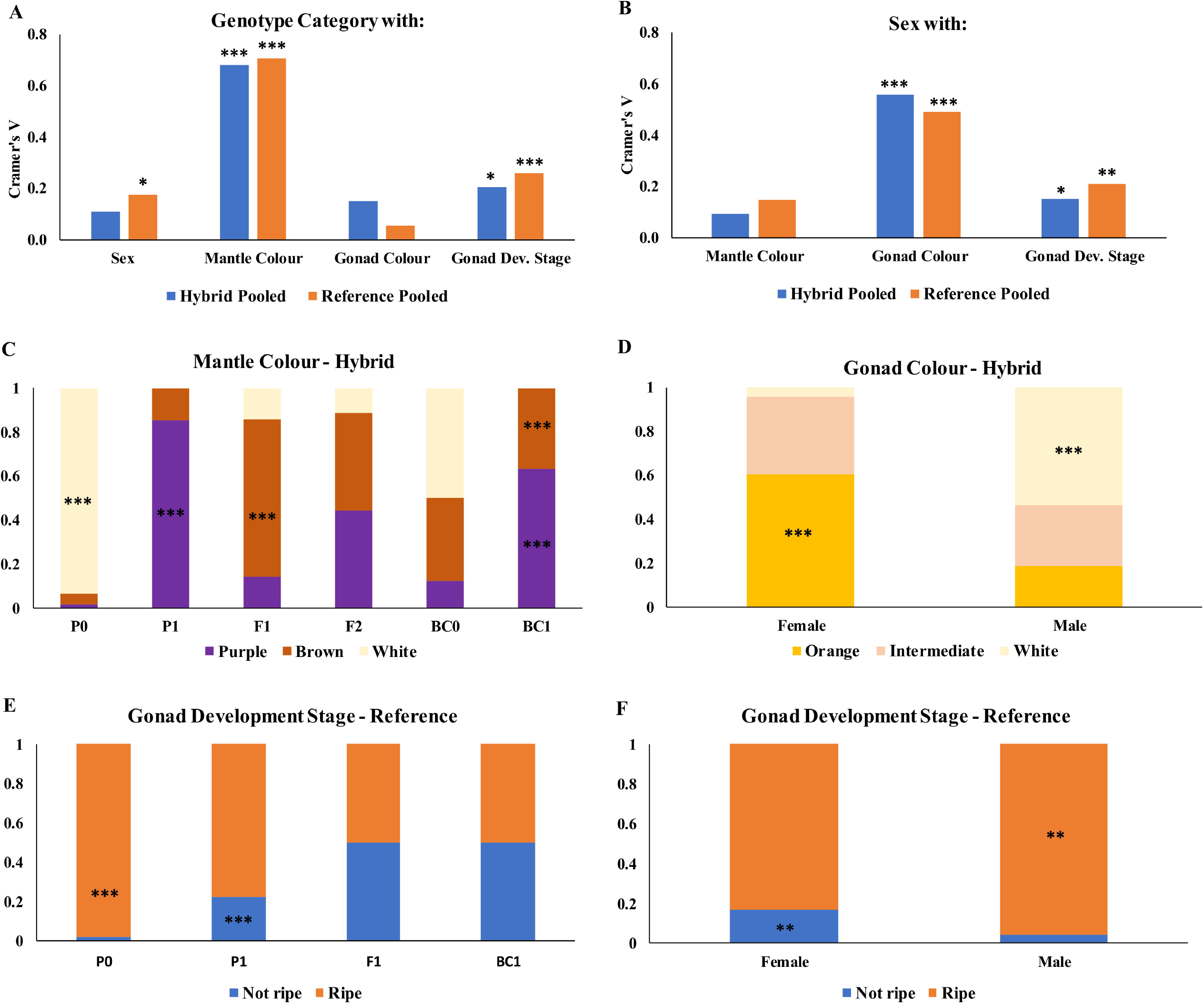
Results of tests of association for pooled reference and pooled hybrid populations. Cramer’s V for association between NewHybrids genotypic categories and phenotypic characters (A) and between Sex and the other phenotypic characters (B). C) to F) distributions for characters as shown on the figure. Significance levels are obtained from adjusted standardised residuals for individual cells in the contingency tables. * P<=0.05;**<=0.01;***<=0.001. Further details are given in **File S4-wsB**.

Mantle colour, gonad colour and gonad development stage show significant differences between populations (**Files S3** and **S4**). There is no variation in sex ratio between populations. In addition, a binomial test against a 1:1 expectation is not significant over all populations pooled (p=0.922). There is a small but significant association between genotypic category and sex for the reference populations (**Figure 7**) with females more frequent in P0 and males in P1 (**File S4-wsB**). These results are also in line with other studies for mussels reporting a 1:1 sex ratio (Seed 1969; Fisher & Skibinski 1990). Although marked sex ratio differences exist between individual crosses (Saavedra et al. 1997; Kenchington et al. 2002) these would be expected to average out in population samples. Thus, the cause of the association between genotypic category and sex for the reference populations is unclear.

There is a strong and significant association of genotypic category with mantle colour as expected (**Figure 7**, panel A). There is also a strong association between gonad colour and sex (panel B). Gonad development stage shows less strong but still significant associations with genotypic category and with sex (panels A and B). The distributions for specific phenotypes for mantle and gonad colour are shown in panels C and D for the hybrid populations pooled with instances of significant excess of observed over expected indicated on the histogram bars. For mantle colour a significant excess of purple is observed for BC1 as well as P1 which is explicable given the high *M. galloprovincialis* ancestry for this genotypic frequency category. Of note is the high and often significant excess of brown mantle colour in the F1 and admixed genotypic frequency categories compared with P0 and P1. For gonad colour there is a significant excess of orange and white in females and males respectively in the hybrid populations. This also occurs in the reference populations. For the reference populations, association between genotypic category and gonad development is towards a significant excess of “ripe” and “not ripe” gonads in P0 and P1 respectively (panel E) and a significant excess of “ripe” and “not ripe” in male and female respectively (panel F).

## Discussion

### Early Admixture Studies

Phenotypic differences distinguishing two distinct forms of mussel in Southwest (SW) England were reported over 50 years ago (Lewis & Seed 1969; Seed 1971). The hypothesis that these comprise *M. edulis* and *M. galloprovincialis* was supported by allozyme studies which confirmed genetic differences between the forms and that the latter resembled Mediterranean *M. galloprovincialis* (Ahmad & Beardmore 1976; Skibinski et al. 1978a,b). The allozyme studies were based on a small number of partially diagnostic loci (Skibinski et al. 1980) and suggested greater admixture on the south than the north coast of SW England (Skibinski et al. 1983).

### Hybrid Zones in North and South of SW England

The studies with the locus Glu-5 provide evidence of a hybrid zone from Start Point to St Ives (see **Figure 1**) (Hilbish et al. 1994; Gilg & Hilbish 2000; Hilbish et al. 2002.; Gilg & Hilbish 2003a). Parental *M. edulis* and *M. galloprovincialis* populations occur to the east of Start Point and to the northeast of St Ives respectively. There is some dispersal from the hybrid zone into the parental populations, but little evidence of dispersal into the zone. There could also be dispersal of *M. edulis* into the hybrid zone from an unidentified source. In addition, self-recruitment, which may be important in mussel populations (Bierne et al 2003b; Gilg et al. 2007; Gosling et al. 2008) could occur within the zone.

Porthcurno, Carlyon and Whitsand are spaced apart within this hybrid zone, whereas Padstow and Bude are within the *M. galloprovincialis* region. Croyde being further northeast is far outside this region. The possibility that dispersal is occurring northeast of St Ives out of the hybrid zone (Gilg & Hilbish 2003a,b) can be tested. At Padstow, Bude and Croyde there are five F1 and two BC1 individuals (**Figure 3**). If these individuals derive by dispersal north from the hybrid zone between St Ives and Start Point they should resemble genetically mussels from this region rather than those north of St Ives. The SNP18 locus provides the possibility of a test as the frequency of the G allele differs markedly between P1 individuals in the north and south coast populations (**Figure 6**). Three groups of mussels are compared, the five F1 and two BC1 individuals (“north hybrid”), the remaining mussels on the north coast (“galloprovincialis”) and mussels in Porthcurno, Carlyon and Whitsand (“south”). Excluding P0 individuals, the E allele proportion for these three groups are 0.857, 0.933 and 0.386 respectively. In pairwise contingency tests the “north hybrid” vs “galloprovincialis” test was not significant (p=0.423), but the “north hybrid” vs “south” comparison was highly significant (p=0.004). This suggests that the hybrid individuals are not immigrants from the southern coast hybrid zone but are produced from a second hybrid zone flanked by *M. galloprovincialis* populations to the south-west and *M. edulis* populations further to the north-east or even in South Wales. An additional test of dispersal northeast of St Ives based on the NewHybrids G3 categories is described in **File S3**. Recently we have genotyped 20 mussels collected in 2018 at Dartmouth east of Start Point for the same SNPs (see **Figure 1** and **File S1-wsO**). Eighteen of the mussels are classified as P0 by NewHybrids confirming that this is an *M. edulis* region. However, one P1 and one BC0 mussel were found (**File S1-wsO**). This contrasts with previous reports that found only *M. edulis* in this region (see Gilg & Hilbish 2003a).

The extent of admixture in hybrid populations can vary depending on the *Mytilus* species involved. The pattern of rising and falling of the G allele frequency from Start Point to the Croyde region is similar to the pattern of changes observed on the Atlantic coast of France from the Bay of Biscay to Normandy (Bierne et al. 2003a). There is also evidence of higher introgression rates in European compared to American *M. edulis* x *M. trossulus* hybrid zones (Fraisse et al. 2016; Simon et al. 2021). In SW England the significant differences in the frequency of the G2 genotypic categories between the hybrid populations (**Figure 3** and **Figure 5**) establish that they are not replicate samples from a homogeneous extended hybrid zone. The results for the SNP18 locus also suggests that the *M. galloprovincialis* individuals on the north and south coasts of SW England have different origins.

### Interbreeding and Production of F1 and F2 Individuals

Though the number of F1 individuals identified is small in the four hybrid populations (9/239 or 3.8%) (**File S2-wsE)** there is a significant excess at Croyde relative to the others (**File S2-wsH**). Segregation of E and G alleles is not involved in the production of F1 from P0 and P1 parents. There can thus be some confidence that F1 individuals are genealogically first-generation hybrids (f1). Croyde has no other admixed individuals identified by NewHybrids (**Figure 3, File S2-wsF** (Table A)). It thus differs markedly from the three other hybrid populations and stands out as a population where interbreeding is at the early stages.

A low frequency of first-generation hybrids (f1) in wild mussels from hybrid zones suggests the possible involvement of prezygotic and both exogenous and endogenous postzygotic reproductive barriers (Wilhelm & Hilbish 1998; Bierne et al. 2006; Diz & Skibinski 2007; Brannock & Hilbish 2010). Since admixed individuals are frequent overall in the hybrid populations, low viability of f1 seems unlikely. A low rate of generation of f1 would be favoured by differences in spawning time between *M. edulis* and *M. galloprovincialis*. Reproductive cycle differences have been observed between sites and seasons but still overlap, with *M. edulis* spawning earlier in the year (Seed 1971; Gardner 1992; Secor et al. 2001; Bierne et al 2003b; Gilg & Hilbish 2003c; Bignell et al. 2008). The higher frequency of “not ripe” mantles in the *M. galloprovincialis* at Padstow and Bude compared with Swansea where “ripe” is dominant (**File S4-wsA**) is consistent with this given that mussels studied here were collected in March. The hybrid populations do not show well-defined spawning periods as “ripe” is significantly more frequent at Porthcurno and “not ripe” more frequent at Whitsand. Some complexities in spawning pattern are evident, for example in the association between sex and gonad development stage (**Figure 7**, panel F, and **File S4-wsB**). A difference in spawning time between sexes was previously observed in *M. galloprovincialis* with males spawning earlier than females, but with synchronous spawning in *M. edulis* (Secor et al. 2001). When values for “ripe” (as a %) are plotted as a line graph from west to east, separately for the north and south coast populations in **File S4-wsE**) there is an interaction such that “ripe” increases from west to east on the north coast but decreases from west to east on the south coast. In regression analysis, the north-south difference is significant (p=0.034) as is the interaction (p=0.017). The cause of this complexity is not immediately obvious. It is surprising that the frequency of F2 is so high (9/239 or 3.8%) given the low frequency of F1 individuals in hybrid populations in SW England (3.8%) and that f2 individuals must be produced from f1 x f1 crosses. A spawning time for f1 intermediate between the parental species would favour a high frequency of such crosses.

### Asymmetrical Introgression

In the triangle plots there are more BC1 individuals on the left hand side than BC0 individuals on the right hand side (**Figure 3**). The greater frequency of BC1 suggests that admixed individuals are backcrossing more often to *M. galloprovincialis* than to *M. edulis*. This would favour higher introgression of *M. edulis* alleles into *M. galloprovincialis*. As BC1 has a significantly higher frequency in the large than small size class, a contributing fact might be higher viability of BC1. The possibility of asymmetric introgression has a parallel in early work on mitochondrial DNA in the hybrid zone. Haplotypes at high frequency in *M. galloprovincialis* from Bude and Padstow were absent from Swansea though haplotypes at high frequency in Swansea were present in Bude and Padstow (Edwards & Skibinski 1987). This would be generally consistent with gene flow from *Mytilus edulis* to *M. galloprovincialis*. Predominant gene flow from *M. edulis* to *M. galloprovincialis* is also consistent with the occurrence of *M. edulis* in SW England but the absence of *M. galloprovincialis* in South Wales. Asymmetric introgression of E alleles into *M. galloprovincialis* has also been observed in French hybrid populations (Fraisse et al. 2014; Simon et al. 2021), as well as into *M. trossulus* in Europe. This might reflect an adaptive advantage of *M. edulis* alleles in other species (Fraisse et al. 2016; Simon et al. 2021).

### Causes of Population Differences

A variety of factors have been identified which might contribute as causes of differences between populations in the SW England hybrid zone. These include factors influencing *M. edulis* and *M. galloprovincialis* differentially such as selective viability (Skibinski 1983; Gardner & Skibinski 1988, 1991a; Skibinski & Roderick 1991; Gardner et al. 1993; Wilhelm and Hilbish 1998; Bierne et al. 2002a) and growth rate (Seed 1971; Gardner & Skibinski 1991b; Gardner et al. 1993; Hilbish et al. 1994). Many of these are also likely to be affected by endogenous and exogenous environmental factors differing between localities (Wilhelm & Hilbish 1998; Gilg & Hilbish 2003a,c; Bierne et al. 2006). To build and test explanatory models of the hybrid zone incorporating these diverse factors will be a substantial undertaking. The possible adaptive significance of the phenotypic characters studied here can be considered. Mantle edge colour is one of the most important distinguishing features between the two species (Lewis & Seed 1969). Darker mantles in *M. galloprovincialis* may be an evolutionary response to more intense sunlight in the Mediterranean (Seed 1971). In the present study there is a marked difference in colour between P0 (white) and P1 (purple) (**Figure 7**). The hybrid categories show a higher frequency of individuals with brown mantles, significant in F1 and BC1 (**Figure 7**, panel C). The excess of brown mantle in the F1 suggests an allelic or non-allelic interaction associated with high heterozygosity. Although early research reported that female and male gonads tend to be pinky-orange and creamy-white respectively (e.g., Seed 1969), more recent work suggested that gonad colour is not a reliable trait to determine sex (Mikhailovic et al. 1995; Petes et al. 2008). The results of the present study support the earlier finding (**Figure 7**, panel D). Because consumers may have a preference for mussels with particular gonad colour, for example orange which is due to carotenoids, the present result may have an important application in aquaculture. For example, mussels with particular gonad colour might be generated in hatcheries with sex biased crosses. A heritability of 0.9 obtained for this character in *M. galloprovincialis* (Pino-Querido et al. 2015) suggests possible modification by conventional selective breeding or strain selection.

### Long Term Stability of Hybrid Zones

In mussels collected in SW England hybrid populations from 1980-81 to 1996-98 the frequency of *M. galloprovincialis* was greater in larger and older than smaller and younger mussels (Skibinski 1983; Gardner & Skibinski 1988; Skibinski & Roderick 1991; Gardner et al. 1993; Wilhelm & Hilbish 1998; Hilbish et al. 2002). Three hypotheses were suggested to explain this, differential growth, differential mortality or viability, and an historical change involving replacement of *M. galloprovincialis* within the hybrid zone (Skibinski 1983; Gardner & Skibinski 1988). Higher mortality of *M. edulis* and viability of *M. galloprovincialis* is the preferred hypothesis (Gardner et al. 1993; Wilhelm & Hilbish 1998). Higher strength of attachment to the substrate or higher resistance to thermal stress favouring *M. galloprovincialis* have been proposed as selective factors (Willis & Skibinski 1992; Hilbish et al. 1994; Hilbish et al. 2002; Riginos & Popovic 2020). The apparent stability of the hybrid zone might be due to the compensating immigration of *M. edulis* (Edwards & Skibinski 1987; Skibinski & Roderick 1991; Wilhelm & Hilbish 1998).

The relationship between shell length and *M. galloprovincialis* allele frequencies has been updated to 2016 incorporating Est-D, Glu-5 and the SNP loci for samples from Whitsand, Croyde, and Porthcurno (**Figure 8**). The positive relationship between shell length and the G allele frequencies is generally similar in these different studies over a period of 35 years. This apparent stability of the hybrid zone can be considered in relation to global warming. Laboratory studies suggest that *M. galloprovincialis* is better able to cope with warmer temperatures than *M. edulis* (Fly & Hilbish 2013). In studies of three hybrid zones of these species in France over a period of two decades, one of the zones changed its position into warmer water (Hilbish et al. 2012), and an increase of *M. galloprovincialis* on the Irish coast was reported during a time interval of 25 years (Gosling et al. 2008). Sea surface temperature (SST) in the UK showed a small gradual increase of 0.2 to 0.3 ^O^C from 1870 to around 1980, but from 1980 to 2020 showed a more dramatic increase of 0.7-0.8 ^O^C. The timing of this large change in SST coincides with the studies of **Figure 8**. Thus these hybrid populations have not responded to warming waters by a shift towards higher G allele frequencies. Introgression in animals might be a source of adaptive genetic variation (Hedrick 2013), related to positive heterotic or overdominant fitness effects in hybrids (Vendrami et al. 2020), which might reduce vulnerability to the effects of climate change through evolutionary rescue (Brauer et al. 2023). Thus, shifts in the ranges of hybridising mussel species in response to global warming might occur in some geographic regions but be resisted in others.

**Figure 8:**
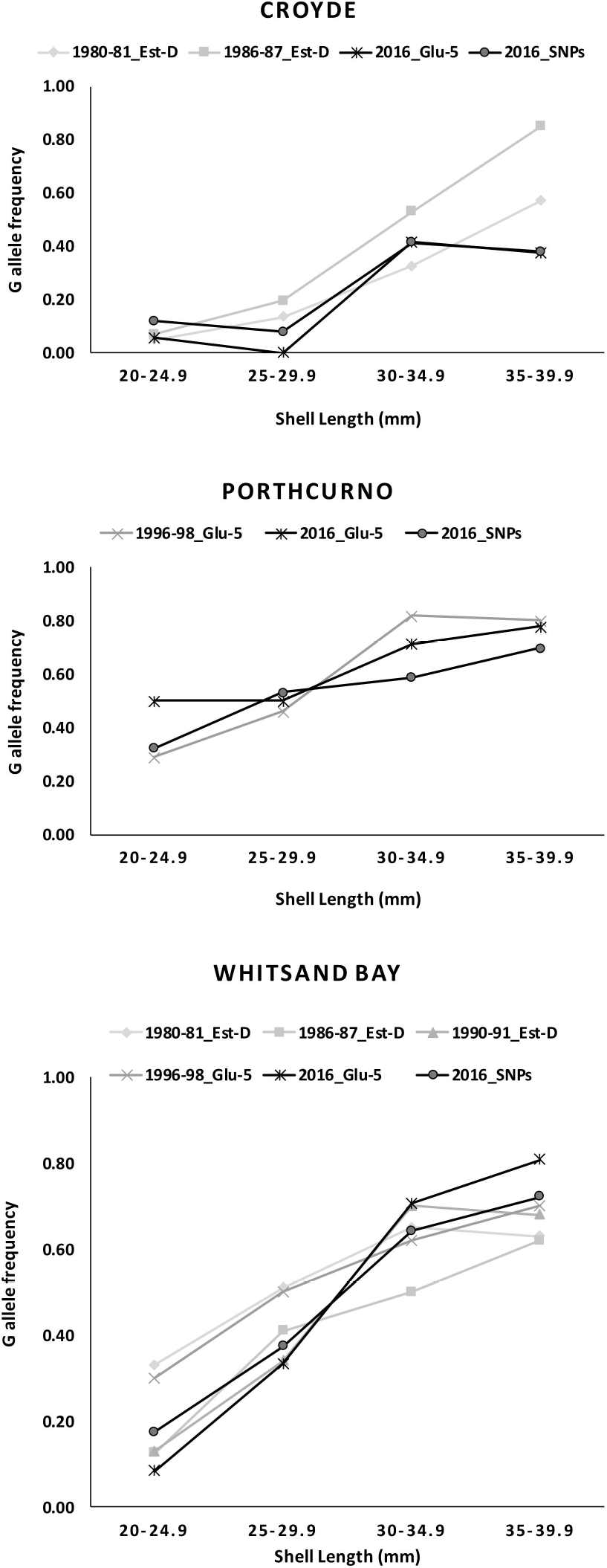
Historic evolution of G allele frequencies obtained from highly ancestry-informative markers in different studies where mussel samples analysed were collected from 1980 to 2016 in the following hybrid populations in SW England: Croyde (CR), Porthcurno (PC) and Whitsand Bay (WB). The molecular markers used (in parenthesis) and studies from which the G allele frequencies were derived are as follows: Skibinski 1983 (*Est-D* allozyme marker; sample collection in 1980-81), Gardner & Skibinski 1988 (*Est-D*; sample collection in 1986-87), Wilhelm & Hilbish 1998 (*Est-D*; sample collection in 1990-91), Hilbish *et al*. 2002 (*Glu5* nuclear marker), and results from the present study deriving from the genotyping of Glu-5 and 57 SNPs (average G allele frequency) in mussel samples collected in 2016.

### Outlier SNP Loci in Hybrid Zones

Outlier loci like SNP18 have been observed in previous studies in France and Spain. Strong introgression was observed at the DAMP2 locus from *M. edulis* into *M. galloprovincialis* (Bierne et al. 2002b). Seven outliers were detected in a study of Atlantic and Mediterranean populations and two of these had high frequencies of *M. edulis* alleles in Mediterranean *M. galloprovincialis* (Gosset & Bierne 2013). An *M. edulis* allele at the mac1 locus also had high frequency in *M. galloprovincialis* possibly attributable to a selective sweep (Fraise et al. 2014). These loci have in common with SNP18 that an *M. edulis* allele has unexpectedly high frequency in *M. galloprovincialis*. Selection acting at a locus may cause it to be statistical outlier (Morin et al. 2004; Lotterhos & Whitlock 2014), thus *M. edulis* alleles might have some selective advantage in *M. galloprovincialis*, perhaps adapting it to colder northern temperatures. A selective effect might be associated with other substitutions or sequences in disequilibrium within the SNP18 polymorphism in the Notch 2 protein. Future work could involve sequencing of copies of the gene and deriving three dimensional structures to try to infer function as has been used for mtDNA proteins (Skibinski et al. 2017). The function of the Notch protein in ectothermic organisms might be influenced by temperature (Shimizu et al. 2014). Thus different alleles could have adaptive significance by promoting optimal development or growth at different seawater temperatures. Evidence in support of the hypothesis that the allele present in *M. galloprovincialis* could have evolved after its evolutionary split from *M. edulis* is supported by the observation that mussel samples from the conspecific *M. trossulus* are fixed for the same allele as in *M. edulis* (Simon et al. 2021; Diz AP, unpublished data), as well as by BlastX analysis (nrNCBI) indicating that Notch sequences from the more distantly related *M. californianus* and *M. coruscus* also have the same allele as in *M. edulis* and *M. trossulus*.

### Extent of Admixture in SW England

The use of many diagnostic SNP loci provides good resolution of the variety of admixed genotypes in the hybrid zone, as evidenced by the diversity of individuals differing in ancestry (Q-value) and individual heterozygosity (Ho) (**Figure 3**). In comparison with the early allozyme studies it also allows analyses of patterns of intercorrelation of many pairs of linked loci (**Figure 4**). An interesting question is whether individual mussels in the triangle plots belong to genealogical classes reflecting few or many generations of interbreeding. This is of relevance to assessing the strength of the barrier to admixture and introgression between the species. *M. edulis* and *M. galloprovincialis* mussels have been recognized as such in SW England over many decades of research and can be equated with the p0 and p1 genealogical classes which in turn are picked out as P0 and P1 by NewHybrids. Each year p0 and p1 mussels as well as f1 will be regenerated. This process might be balanced by the loss of more highly admixed individuals and their lineages from the zone, either by drift or selection. There might be some confidence that the G2 analysis picks out individuals in the specific genealogical classes, for example that F2 individuals are f2. Many individuals assigned as P1 in the NewHybrids G2 analysis are reassigned as genotypic categories unique to the G3 analysis such as P1_BC1 (**File S2-wsE**). Whether these would belong to the genealogical class p1_bc1 is less certain.

The estimates of the number of generations since admixture from STRUCTURE range between 0.2 generations at Croyde and 4.3 generations at Whitsand with Porthcurno (2.0) and Carlyon (2.3) intermediate. This is consistent with the hypothesis that many of the admixed individuals derive from only a small number of generations of ongoing interbreeding. If the admixed individuals had been the result of many generations of interbreeding, recombination might have substantially reduced R1 and R2 (**Figure 4**). The contrast between Croyde and Whitsand in the values of R1 and R2, lower in the latter is consistent with the estimated difference in generations since admixture. However, for Porthcurno and Carlyon which have intermediate values the pattern is less clear. Porthcurno appears to resemble Whitsand in R1 and R2 and Carlyon is perhaps closer to Croyde.

The results for generations since admixture contrast with the results from dock populations where complete mixing of gene pools might be expected and where estimates of number of generations since admixture are higher ranging from 4-14 (Simon et al. 2020). Techniques for interbreeding mussels in the laboratory are improving (Kenchington et al. 2020). Thus if a high density linkage map can be obtained for *Mytilus* in future, better information on genealogical classes and the ancestral relationship between individual mussels should be accessible as in human genetics (Henn et al. 2012). Such a linkage map would also allow great progress towards determining the genetic basis and adaptive significance of variation of mantle colour, gonad colour and other phenotypic attributes through genome wide association studies (Uffelman et al. 2021; Peñaloza et al. 2022; Luo et al. 2023).

## Acknowledgments

Nerea González-Lavín, Juan Pablo Pérez-Diz and the marine station staff (ECIMAT, CIM-UVigo) for their indispensable assistance in sampling, sample processing, and molecular analysis. Alexis Simon and Nicolas Bierne provided valuable advice on SNP development and selection and data analysis and offered constructive criticism on earlier versions of this manuscript. This work was supported by the Spanish “*Agencia Estatal de Investigacón (AEI)*” (codes AGL2014-52062-R and PID2019-107611RB-I00), Fondos Feder (ERDF, European Commission), and Xunta de Galicia (“*Grupos de Referencia Competitiva*” ED431C 2020/05).

## Data Accessibility and Benefit-Sharing

Raw data underlying the main results of the present study are provided in **File S1**. Benefits from this research accrue from the sharing of our data and results of present work in an Open Access journal.

## Author Contributions

**APD**: Conceptualization, Methodology, Investigation, Formal analysis, Writing – original draft, review & editing, Funding acquisition. **DOFS**: Conceptualization, Methodology, Investigation, Formal analysis, Writing – original draft, review & editing.

