## Supplementary material for "Spatial and long-term temporal evolution of a marine mussel hybrid zone (*Mytilus* spp.) in SW England": File S3

**NewHybrids analysis**

The analysis described in the main text is for two generations of interbreeding (G2 analysis). Some additional methodological details and results are described here. The NewHybrids method has previously been applied to analyse up to three generations of interbreeding (G3 analysis) (Pritchard et al. 2016; Chhatre et al. 2018; Taillebois et al. 2019; Buck et al. 2020), and this is also explored.

In the main analysis described in the text additional mussels were included solely to increase the power of the analyses on the main sample of 480 mussels. Information on these mussels is shown in **File S1-worksheet(ws) E**. These mussels were of two kinds. Some were used in separate spawning experiments (from row 485). The remainder were from additional non-UK reference populations for *M. edulis* and *M. galloprovincialis* reported by Simon et al. (2020) (from row 622). The results and analysis reported in the main text are for the 60 mussels for each of the 8 populations and exclude these additional mussels. In what follows below this is referred to as the "all mussels" dataset and this was used for the analysis reported in the main text.

In addition, the above analysis was repeated excluding the population sample from Vigo, additional spawned mussels from Vigo and the additional reference populations. For these analyses the results reported are for the 60 mussels from the 7 UK populations. This is referred to as the "UK mussels" dataset.

The NewHybrids method was developed primarily for unlinked markers (Anderson & Thompson 2002). In most studies of natural populations, it would have been applied to datasets where linkage information is unavailable, with the hope that the method would be robust to deviations from this assumption. A third analyses was carried out using the "all mussels" dataset but using only 15 unlinked SNPs according to the genetic map available (**File S2-wsA**). This is referred to as the "15 loci" dataset. In all the NewHybrids runs, Jeffreys-like priors for both mixing proportions and allele frequencies, and 100,000 burn in and 1,000,000 iterations were used.

With three generations of interbreeding (G3) there are 21 different genotypic frequency categories. However, some of these have identical values for the array of the expected genotypic proportions. For example, proportions of 0.25 for each of EE, EG, GE and GG occur with F2 progeny, and with progeny of the F2 x F2 and F1 x F2 crosses. These are pooled and treated as a single category. There are 15 unique G3 categories having different genotypic proportions (**File S2-wsB)**. Some of these have identical heritage values for the proportion of the G (*M. galloprovincialis*) allele, for example F1 and F2. Some of the G3 categories correspond to a single, others to more than one genealogical class. Three are from G2 (e.g., P0) three are mixed G2 and G3 (e.g., BC0 with P0_F2), seven are unique to G3 (e.g., P1_BC1) and two are mixed G3 (e.g., F1_BC0 with F2_BC0) (**File S2-wsB**). If interbreeding had proceeded only to the second generation, a G3 analysis would have the possibility to pick out the six genotypic categories that contain those from G2. If the G3 analysis does assign many individuals to the unique or mixed G3 categories, it could be possible to argue that interbreeding has proceeded to at least the third generation. It is unlikely that a G4 analysis would be useful because the different genotypic frequency categories become too many and too similar in expected genotypic proportions for discrimination given available sample sizes.

For the G2 analysis for the "all mussels" dataset the genotypic category assignations for individual mussels for each population are given in **File S2-wsD**, with numbers summarised in **wsC**. In each population, mussels are ranked in size from small to large along the X axis. Even though the samples from the reference populations Swansea (P0, *M. edulis*) and Vigo (P1, *M. galloprovincialis*) are not fixed differently at the SNP loci (**Figure 2** and **File S1**) all the mussels are correctly assigned by NewHybrids. In the Bude reference population all individuals are correctly assigned as P1 except one which is assigned with high probability as a backcross to *M. galloprovincialis* (BC1). In Padstow all except three individuals are assigned as P1. These exceptions are assigned with high probability, one as BC1 and two as F1.

The other four populations Croyde, Porthcurno, Carlyon, and Whitsand show evidence of mixing of the two species as well as interbreeding and can be regarded as hybrid populations. Visual examination of the histogram suggests that the genetic structure is different between these populations. Thus, at Croyde there is mixing but little admixture beyond the F1, whereas in the populations to the south there is substantial admixture with many individuals assigned as backcrosses to *M. galloprovincialis* (BC1) particularly at Porthcurno and Whitsand. The contrast between the large and small mussels at Croyde is very evident in **File S2-wsD**, as most of the individuals not assigned as *M. edulis* (P0) are seen to be larger mussels to the right of the histogram. Individuals were assigned to the genotypic category for which they had the highest posterior probability, and a number of statistical comparisons were made using contingency analysis as described in the main text of the paper.

Results for the NewHybrids analysis taking account of three generations of interbreeding (G3) are shown for individual mussels for each population in **File S2-wsD**, with numbers summarised in **wsC**. The contrast with the G2 results is evident. Many individuals have been reassigned, for example many P1 individuals in the reference populations Bude and Padstow.

These differences have been summarised and quantified for the reference populations pooled and hybrid populations pooled in **File S2-wsE**. This shows the number of individuals in each of the six G2 categories which become assigned to the various G3 categories. Not all of the 15 unique G3 categories (**File S2-wsB**) have been observed in the data. Some individuals are assigned the same categories in G3 as they are in G2, for example many P0 and P1 individuals and also five of nine F1 individuals, but many are reassigned as new categories. This result is consistent with the hypothesis that interbreeding in the hybrid zone is proceeding beyond the second generation. Of the 165 P0 individuals for G2 (105 from the hybrid populations and 60 from the reference populations), 162 are reassigned as P0 the other three as P0_BC0 for G3. This is in clear contrast to what is observed for P1. In the hybrid populations, of 48 individuals classified as P1 for G2, 33 are reassigned as P1_BC1 or BC1_BC1 for G3. Even in the reference populations, 19 of 176 P1 individuals for G2 are reassigned as P1_BC1 for G3. This marked asymmetry is consistent with greater backcrossing to *M. galloprovincialis* (P1) than to *M. edulis* (P0) within the hybrid zone, and is in agreement with other evidence presented in the main text. Consistent with this asymmetry is the observation that 54 of 56 BC1 individuals are reassigned as BC1_BC1 or P1_BC1 as described above. In the main text of the paper it is noted that in the G2 analysis 9/239 = 3.8 % individuals are classified as F2 which is surprising given that the frequency of F1 is low. One explanation arising from the G3 analysis is that many of these F2 classifications are incorrect. This is because all of these nine F2 individuals have been reassigned as categories occurring only in G3 despite F2 being one of the options in the G3 analysis.

A breakdown by population of the data of **File S2-wsE** is given in **File S2-wsF**. At Croyde, the three F1 individuals identified in the G2 analysis are reassigned as F1 in the G3 analysis, consistent with their arising directly from P0 x P1 crosses. However, two P0 and two P1 individuals are reassigned as P0_BC0 and P1_BC1 respectively. These individuals require prior backcrosses with F1 to give BC0 and BC1 respectively. Although individuals of the latter two genotypic categories were not detected at Croyde, numbers are small, and this sequence of crosses seem feasible at Croyde. At Bude eight of 51 P1 individuals are reassigned as P1_BC1 and one BC1 individual as BC1_BC1. At Padstow eleven of 46 P1 individuals are classified as

P1_BC1, one of two F1 individuals is reassigned as BC0_BC1 and a single BC1 individual as P1_BC1.

A feature of the crosses leading to the reassignations in these north Cornwall populations north east of St Ives is that they feature P1 predominantly. However, P0 must also be involved to generate F1 prior to backcrossing. The hypothesis that dispersal from the hybrid zone beyond St Ives into north Cornwall (Gilg & Hilbish 2003a,b) is not well supported. It seems more likely that these individuals are produced from a second hybrid zone in the Croyde region as described in the main text. In this circumstance the exceptional individuals at Bude and Padstow may derive from this second hybrid zone. This is backed up by other analysis described in the main text. An additional test of dispersal northeast of St Ives based on the NewHybrid genotypic frequency categories for three generations of interbreeding (G3) with results set out in **File S2-wsF** (Table A). Of 128 P1 individuals at Padstow, Bude and Croyde, 21 were reassigned to the new category P1_BC1. It is not suggested that these necessarily belong to the genealogical class that are offspring of the cross p1 x bc1. Nevertheless, they might be amongst the P1 mussels having the greatest *M. edulis* ancestry. When pooled these 21 P1_BC1 mussels have a frequency of the SNP18 E allele of 0.909, which is in line with the overall high frequency shown in **Figure 6** for P1 in these northern populations. Had they originated through dispersal from the region between St Ives and Start Point a lower frequency of the E allele would be expected (**Figure 6**). A previous study of mtDNA variation also supports a substantial genetic difference between Whitsand and Croyde (Edwards & Skibinski 1987). Whitsand in common with *M. edulis* from Swansea lacked a haplotype that was common at Padstow, Bude and Croyde.

The assignation of individuals to genotypic categories by NewHybrids will depend in part on the populations included in the analysis. The *M. galloprovincialis* individuals in the UK might have a higher level of *M. edulis* ancestry than those in Vigo which are geographically more distant from *M. edulis* populations. Thus, when Vigo is included, UK individuals assigned as P1 in the G2 analysis, and having more *M. edulis* ancestry, might more readily be assigned as BC1, P1_BC1 or BC1_BC1. When Vigo is excluded, such individuals might more readily be assigned as P1. To test this, the NewHybrids analysis was repeated with the "UK mussels" dataset. The results are compared for both analyses in **File S2-wsE** with the results broken down by population in **File S2-wsF**. The results do indicate that for the hybrid populations, fewer P1 individuals are assigned as BC1 in the G2 analysis, 37 compared with 54 when using the "UK mussels" dataset. However, many BC1 mussels are still reassigned as P1_BC0, P1_BC1 or BC1_BC1 whether or not UK populations are included. Thus reassignation of P1 mussels in the G3 analysis cannot be said to be an artefact of including mussels from outside the UK.

The G2 analysis was repeated for the "15 loci" dataset. The results are presented in **File S2-wsG** and compared with the results for the "all mussels" dataset. Tables give the numbers of individuals in each population and genotypic frequency category aligned for the two datasets. The correlation between the two datasets is very high (0.993). We thus decided to base our main analysis on the use of all 57 SNPs to benefit from the information from all the loci available at the cost, in relation to the assumptions of the NewHybrids method, of using some loci that are linked.

**Phenotypic Character Associations**

The differences between populations in the distribution of sex, mantle edge and gonad colour, and gonad development stage will be largely a reflection of their differences in *M. edulis* and *M. galloprovincialis* ancestry. These population distributions are given in **File S4-wsA** as tables and histograms, together with table-wide p-values for Fisher's exact test together with Cramer's V and information on which individual cells have a significant excess of observed over expected according to the adjusted residual which is converted into a p-value. A summary of the results (**Figure 7**) is considered in the main text and is also referred to below.

As expected, there are large differences in the distribution of mantle colour between populations overall with white being associated with *M. edulis* and purple with *M. galloprovincialis* (**File S4-wsA,** p=0.000). In the reference populations outside the hybrid zone, this pattern is followed with all Vigo individuals having purple mantles and all Swansea individuals having white mantles. The brown mantle colour does appear from the histogram to be more frequent in the four hybrid populations and is highly significant at Porthcurno (p=0.000). Gonad colour also varies significantly between populations (p=0.008) with orange being more frequent in Padstow (p=0.003) and Whitsand (p=0.004), and white slightly more frequent at Bude (0.032), but there is no obvious pattern separating the hybrid and reference populations. The distribution of gonad development stage also varies significantly between populations (p=0.000), with not ripe gonads in excess at Bude (p=0.001), Padstow (p=0.000) and Whitsand (p=0.040), and ripe gonads in excess at Vigo (p=0.000), Swansea (0.001) and Porthcurno (p=0.001). There is no obvious pattern separating *M. edulis* and *M. galloprovincialis* reference populations and the hybrid populations.

The relationship between the genotypic categories and sex, mantle and gonad colour, and gonad development has been analysed. Sex also has some considerable theoretical importance in relation to the phenomenon of doubly uniparental inheritance of mitochondrial DNA in *Mytilus* (Fisher & Skibinski 1990; Skibinski et al. 1994a,b; Zouros et al. 1994a,b). Therefore, the relationship between sex and mantle and gonad colour, and gonad development has also been further analysed.

These two groups of analyses were carried out for the hybrid populations pooled (Croyde, Porthcurno, Carlyon, and Whitsand) and separately for the reference populations pooled (Vigo, Swansea, Bude and Padstow). Cramer's V are plotted for the association tests together with significance levels indicated in **Figure 7** (panels A and B) with underlying statistics in **File S4-wsB**). The *M. edulis* and *M. galloprovincialis* individuals with their distinctive phenotype differences define the P0 and P1 categories in the NewHybrids analysis and completely dominate numerically in the reference populations. Since these categories are also at high frequency in the hybrid populations it is not surprising that there is good correspondence in the values of Cramer's V and significance levels between the reference and hybrid populations in **Figure 7** (panels A and B). The highest associations are between genotypic category and mantle colour and between sex and gonad colour, and there is a lower but still significant association between gonad development stage and both genotypic category and sex.

The distributions of these characters in the contingency tables are shown in **Figure 7** (panels C to F) with instances of significant excess of observed over expected, determined from adjusted residuals, indicated on the histogram bars. Underlying statistics are given in **File S4-wsB**, and histograms for all the contingency tests, with associated tables given in the same file ws**C** and ws**D**. There is no significant difference in sex ratio between categories for the hybrid populations pooled (p=0.815). However, a significant difference was observed for the reference populations pooled (p=0.038) with an excess of females in P0 and males in P1 (**Figure 7**, panel A, and **File S4-wsB** and **wsC**).

In the hybrids there is a highly significant excess of individuals with white and purple mantles in the P0 and P1 categories respectively (**Figure 7**, panel C, **File S4-wsB**). Brown mantle is at higher frequency in the first (F1) and second-generation hybrids (F2, BC0, and BC1), if not significant, but is highly significant for F1 and BC1. There is also a highly significant excess for purple mantle in BC1. In the hybrids there is a highly significant excess of orange and white gonads in females and males respectively (**Figure 7**, panel D), which occurs also in the reference populations (**File S4-wsB**). For the reference populations, association between genotypic category and gonad development is towards a significant excess of ripe and not ripe gonads in P0 and P1 respectively (**Figure 7**, panel E, **File S4-wsB**) and a significant excess of ripe and not ripe in male and female respectively (**Figure 7**, panel F). The trend is in the same direction but not so marked in the hybrid populations (**File S4-wsB** and **wsD**).

### Cited References


Anderson, E. C., & Thompson, E. (2002). A model-based method for identifying species hybrids using multilocus genetic data. *Genetics*, *160*(3), 1217-1229.

Buck, R., Hyasat, S., Hossfeld, A., & Flores-Rentería, L. (2020). Patterns of hybridization and cryptic introgression among one- and four-needled pinyon pines. Annals of botany, 126(3), 401–411.

Chhatre, V. E., Evans, L. M., DiFazio, S. P., & Keller, S. R. (2018). Adaptive introgression and maintenance of a trispecies hybrid complex in range-edge populations of *Populus*. *Molecular Ecology, 27(23)*, 4820–4838.

Edwards, C. A., & Skibinski, D. O. F. (1987). Genetic variation of mitochondrial DNA in mussel (*Mytilus edulis* and *M. galloprovincialis*) populations from South West England and South Wales. *Marine Biology*, 94, 547-556.

Fisher, C., & Skibinski, D. O. F. (1990). Sex-biased mitochondrial DNA heteroplasmy in the marine mussel *Mytilus*. *Proceedings of the Royal Society of London. Series B: Biological Sciences*, *242*(1305), 149-156.

Gilg, M. R., & Hilbish, T. J. (2003a). Patterns of larval dispersal and their effect on the maintenance of a blue mussel hybrid zone in southwestern England. *Evolution*, *57*(5), 1061-1077.

Gilg, M. R., & Hilbish, T. J. (2003b). The geography of marine larval dispersal: coupling genetics with fine-scale physical oceanography. *Ecology*, *84*(11), 2989-2998.

Pritchard, V. L., Erkinaro, J., Kent, M. P., Niemelä, E., Orell, P., Lien, S., & Primmer, C. R. (2016). Single nucleotide polymorphisms to discriminate different classes of hybrid between wild Atlantic salmon and aquaculture escapees. *Evolutionary Applications*, *9*(8), 1017-1031.

Simon, A., Arbiol, C., Nielsen, E. E., Couteau, J., Sussarellu, R., Burgeot, T., ... & Bierne, N. (2020). Replicated anthropogenic hybridisations reveal parallel patterns of admixture in marine mussels. *Evolutionary Applications*, *13*(3), 575-599.

Skibinski, D. O. F., Gallagher, C., & Beynon, C. (1994a). Sex-limited mitochondrial DNA transmission in the marine mussel *Mytilus edulis*. *Genetics*, *138*(3), 801-809.

Skibinski, D. O. F., Gallagher, C., & Beynon, C. M. (1994b). Mitochondrial DNA inheritance. *Nature, 368(6474),* 817–818.

Taillebois, L., Sabatino, S., Manicki, A., Daverat, F., Nachón, D. J., & Lepais, O. (2019). Variable outcomes of hybridization between declining *Alosa alosa* and *Alosa fallax*. *Evolutionary applications, 13(4)*, 636–651.

Zouros, E., Ball, A. O., Saavedra, C., & Freeman, K. R. (1994a). An unusual type of mitochondrial DNA inheritance in the blue mussel *Mytilus*. *Proceedings of the National Academy of Sciences of the United States of America, 91(16)*, 7463–7467.

Zouros, E., Ball, A. O., Saavedra, C., & Freeman, K. R. (1994b). Mitochondrial DNA inheritance. *Nature*, *368*(6474), 818-818.
